# New Approaches to Fast Plate Composition, with a Software Implementation for Taxonomy

**DOI:** 10.64898/2026.07.07.736512

**Authors:** Alexandre P Aguiar

## Abstract

The preparation of multi-panel figures remains a labor-intensive step in scientific publication, particularly in image-intensive fields such as zoological taxonomy. Although many tools are available, they are generally either general-purpose applications or highly specialized solutions that may be difficult to install or time-consuming to learn. Here, new approaches are proposed to simplify and formalize multi-panel figure composition, emphasizing rapid layout construction and experimentation while maintaining explicit control over publication parameters. These include spatial inference of intended layouts from the distribution of images on a canvas, editable formulas for representing and modifying layout structure, integrated operations for image arrangement and transformation, and automated procedures for figure numbering, scale-bar calculation, and legend generation. The approaches are implemented and tested in Griphus, a standalone graphical application developed by and for zoological taxonomists. User-defined parameters such as target print dimensions, resolution, spacing, and color mode are maintained throughout the work, while layouts can be interactively constructed, modified, rotated, and regenerated. The resulting composite figure is accompanied by a readable representation of its layout structure and a .gri session file storing images, transformations, and parameters for exact regeneration. Griphus thus provides a practical implementation of the proposed approaches and a complementary environment to professional image-editing software for efficient, reproducible, and exploratory construction of publication-quality multi-panel figures.

## INTRODUCTION

Despite nearly three centuries of research, huge numbers of the Earth’s species remain undescribed. Yet, regardless of how they are described, taxonomy shares with other sciences a persistent bottleneck: formatting research for publication. The preparation of illustrations is often a particularly laborious part of this process, especially given their importance to scientific communication. Figures convey complex information efficiently and enable direct comparison between observations (Tufte 2001; Rougier et al. 2014). In many biological disciplines, multiple images are commonly combined into composite figures to document structures or illustrate variation.

In nineteenth- and early twentieth-century scientific publications, illustrations were commonly organized into plates, full pages of illustrations printed separately from the main text. The term originated from the engraved metal or photographic plates used in printing and early photography (Riding and Head 2019). Modern journals usually refer to plates simply as figures, but the term remains widely used for multi-panel illustrations labeled sequentially (e.g., “Figures 3A–F”). Over time, biological illustration evolved from individualized artistic compositions (see Czapla 2025) toward more standardized layouts designed to facilitate comparisons between structures and taxa.

The effectiveness of such figures, however, depends not only on image quality but also on the clarity and consistency of their visual arrangement. Preparing publication-quality figures is nevertheless a nontrivial task, with best results usually requiring great care and the use of a wide range of digital tools (e.g., Bevilaqua 2020). Limited formal training in visual communication among researchers further contributes to persistent problems in figure design and composition (Rayaroth et al. 2025), while entire disciplines such as graphic semiology (Bertin 1967, 1977) remain largely overlooked in the biological sciences.

Contemporary journal author guidelines reflect the importance of such standardization, with common requirements including sequential panel labeling, consistent lettering, specified raster formats (e.g., TIFF, PNG, JPG), and defined resolution standards. More specific requirements may include exact figure dimensions, column widths, mandatory scale bars, minimal surrounding whitespace, and others.

Despite this standardization, the preparation of multi-panel figures remains a largely personal and manual activity. Researchers typically rely on general-purpose graphics editors or presentation software to arrange images. Although flexible, these tools are not specifically designed for multi-panel composition, particularly not for the requirements of taxonomic work. Two main problems arise as a result: first, using general-purpose tools demands a mindset that is not optimized for plate-building; and second, their use commonly involves repeated and inefficient manual operations such as resizing, cropping, alignment, spacing adjustment, figure numbering, and the calculation, design, and positioning of scale bars.

The aim of this work is to propose and implement new, more efficient approaches to multi-panel figure composition, both conceptual and practical, with particular emphasis on the requirements of taxonomic research. These approaches are implemented in Griphus, a new software application developed to enable their practical application and testing. The software’s operational principles and main functionalities are presented to show how the proposed approaches can facilitate figure production and systematic experimentation with alternative layouts.

## METHODS

All the ideas, concepts, and approaches underlying Griphus originated in a private Python tool developed for the author’s personal use, and were refined over many years. Despite sustained use, the software remained private largely because substantial effort would have been required to restructure it for broader distribution. To overcome this limitation, OpenAI *Codex* was used for systematic refactoring and expansion of the original program and modules, initially consisting of over six thousand lines. Even so, underperforming sections often required separate development and improvement before being incorporated into the main codebase. Producing a stable, distributable version required hundreds of documented hours of development, testing, correction, and redesign. Nearly all frontend artwork is original and similarly required considerable effort and repeated refinement. The software is implemented and maintained as a Python application but is currently distributed only as a standalone compiled executable (.exe). The interface is built with Tkinter, with optional drag-and-drop support via tkinterdnd2. Image processing and transformation rely on Pillow, OpenCV, and NumPy, which provide robust support for raster image manipulation and numerical computation.

## RESULTS

This section is not intended as a user manual for the software; a comprehensive manual is available elsewhere. Instead, it provides a formal presentation of the original concepts and approaches, as well as the original practical solutions proposed to translate them into functional operations. Both are described as an integrated framework, as directly implemented in Griphus. Accordingly, the results focus primarily on original contributions, addressing more conventional functions only when necessary for clarity or to document the implementation of essential basic features.

### Input and Output

Common raster image formats, including GIF, JPG, PNG, and TIFF are, by far, the most widely used formats in scientific publications, and are therefore the main focus of this work. They are supported both as input and output in Griphus, but the software generates four coordinated products: (1) a final composite image rendered at user specified physical dimensions and resolution (DPI); (2) a Formula.txt file containing the symbolic layout formula and panel mapping; (3) an optional Legends.txt file with the complete numbered legends for the images on the canvas and, if present, the respective descriptions of the scale bars; and (4) a .gri session file storing all transformations, numbering settings, image parameters, and the computed layout, allowing reopening, modification, and exact regeneration of the session.

### Improved Operational Logic for Plate Construction

Perhaps the most important requirement for efficient plate construction is a fast and objective method for defining and applying any intended layout structure. Two complementary approaches, interactive and symbolic, provide particularly precise and intuitive solutions, and can be considerably more effective than, for example, attempting to describe a complex layout in an LLM prompt. In the interactive approach, images are distributed approximately on the canvas according to the intended arrangement (as in Figure 1), and an algorithm infers the layout structure from their proximity and relative alignment before constructing the final plate. Alternatively, a symbolic layout formula can be specified or edited to define the grouping structure explicitly. For syntax, a simple option is to use a comma to indicate horizontal grouping, a forward slash for vertical stacking, and parentheses to define precedence. For example, **(1**,**2)/3** specifies two images arranged horizontally above a third image. This notation enables the construction of complex layouts and accommodates most image combinations required for scientific multi panel composition. All these approaches are implemented in Griphus.

**FIGURE 1.**
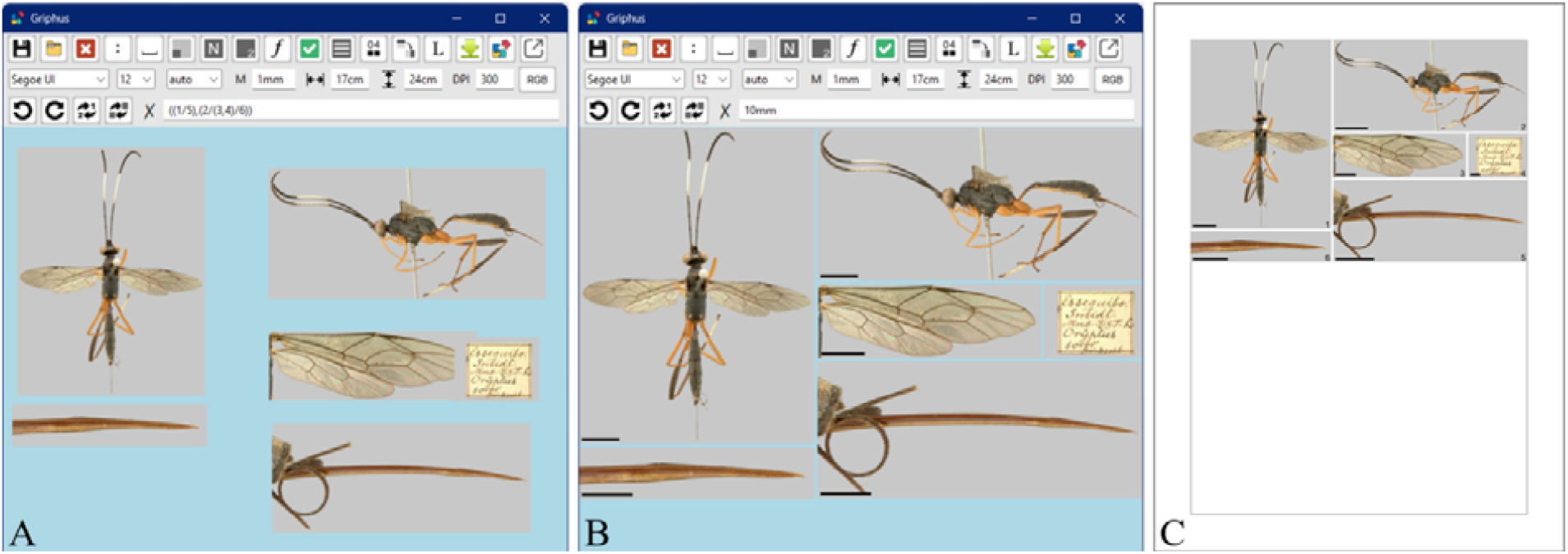
Basic operational logic of the software demonstrated with images of the holotype of *Debilos soror* (Trentepohl, 1829) (Hymenoptera, Ichneumonidae, Cryptinae). Photographs by Berthil B. Longo. **A**, Images loaded by drag-and-drop and manually arranged into the intended groups. The inferred layout formula **((1/5)**,**(2/(3**,**4)/6))**, is calculated and displayed in real time. The formula is fully editable and is needed as a command for multi panel construction. **B**, Approximate canvas rendering of the final multi panel image generated with button 10 (checkmark, green). Scale bars inserted with button 5. **C**, Final image positioned within the target page layout, with the maximum printable area indicated in gray. The image was generated with button 15 (green down arrow) and exported according to the specified parameters, with font sizes calculated to match the target (published) typographic size according to the final image dimensions and resolution.

Panel assembly can also proceed more efficiently when the entire session is governed from the outset by (1) the target journal’s specified physical figure dimensions, expressed in the required units (centimeters, millimeters, or inches), and (2) fixed, user-defined spacing between images. Automatic panel numbering is similarly useful, as it allows labels to be checked and edited while experimenting with alternative layouts. The combined effectiveness of these approaches can be evaluated in Griphus, which incorporates the specified parameters into all layout calculations in real time throughout the session.

Direct access to target plate width, height, resolution (DPI), and color mode (RGB, RGBA, CMYK) as primary interface parameters is particularly important because these values define the physical and digital specifications required for publication. In Griphus, they remain visible and editable throughout the session, while all conversions among physical dimensions, pixel dimensions, and resolution are handled automatically, eliminating repetitive calculations and reducing errors. For TIFF output, the corresponding resolution metadata are also embedded automatically, ensuring that compatible software can correctly interpret the intended physical dimensions of the figure, an important technical requirement that is easily overlooked.

Automatic generation of formatted figure legends from image filenames and corresponding panel numbers can further reduce repetitive work and minimize inconsistencies. When scale bars are inserted (Figure 2), automatically incorporating their values and units into the corresponding descriptions can also be particularly useful. Griphus implements both functions and generates legends in either a full mode, retaining all relevant information from the filenames, or a concise mode, producing shorter descriptions by eliminating repeated terms.

**FIGURE 2.**
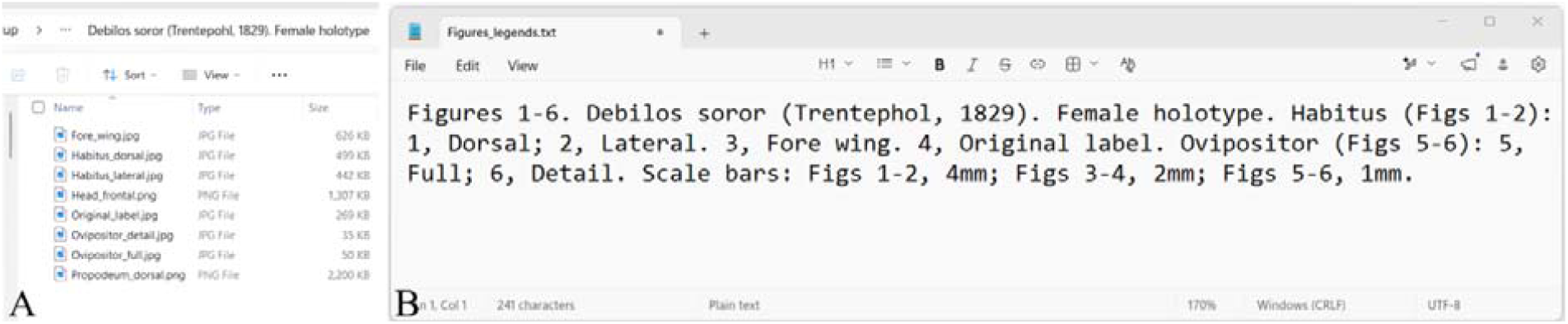
Automatic generation of figure legends, including scale bar descriptions. **A**, Source image files stored in a folder labeled with the species name and type status. **B**, Automatically generated text file containing the legends and scale bar descriptions, opened for editing. Only the images included in the final figure were used to compose the legends (Head and Propodeum excluded).

### Spacing with Strict Aspect Ratio

A central but technically complex feature of plate construction is the assembly of images with constant user-defined spacing, without cropping and while strictly preserving image aspect ratios (as in Figure 1C), regardless of layout complexity. Achieving equivalent results manually is algebraically cumbersome and time-consuming, particularly as the number of images increases or layouts incorporate hierarchical or asymmetric structures, such as **((1**,**2)/3)** or **((1**,**2)/3)/((4/(5**,**6))**, etc. One key advantage, therefore, would be to have these proportions automatically calculated. This is achieved by Griphus.

### Interface Structure and Specialized Functions

A key approach to accelerating plate construction is an interface designed for direct manipulation, with minimal reliance on menus and, where menus are unavoidable, shallow navigational paths. Griphus exemplifies this with only three menus, none containing submenus, and an interface that operates primarily through buttons which activate functions specific to multi-panel composition. Images can be imported conventionally or loaded by drag-and-drop. More routine operations are instead accessible through keyboard and mouse shortcuts while supporting bulk manipulation: any number of selected images can be resized with the mouse wheel, rotated with the arrow keys, and cropped, restored, or flipped using CTRL, ALT, or SHIFT plus the arrow keys, respectively. Although the Delete key is standard for removing images, a triple-click proved more practical and convenient; both methods are implemented in Griphus. Established selection conventions are retained: individual panels are selected with a mouse click, while multiple panels can be selected using CTRL plus click or click-and-drag selection.

Since many plates consist of images arranged in simple rows or columns, these layouts are obvious and intuitive targets for automation, which can substantially accelerate plate construction. In Griphus, this approach can be tested by loading or rearranging entire sets of images at once as rows, columns, proportional rows that preserve relative image dimensions for inspection, or through an original instant-composition procedure (Figure 3A). Instantaneous rotation of the entire layout, without the need to select or reposition individual images, can similarly provide rapid access to alternative arrangements and is also available for experimentation in Griphus (Figure 3B). At the same time, many specialized functions can be devised to optimize specific aspects of multi-panel composition. A range of such functions has been implemented in Griphus and is described below.

**FIGURE 3.**
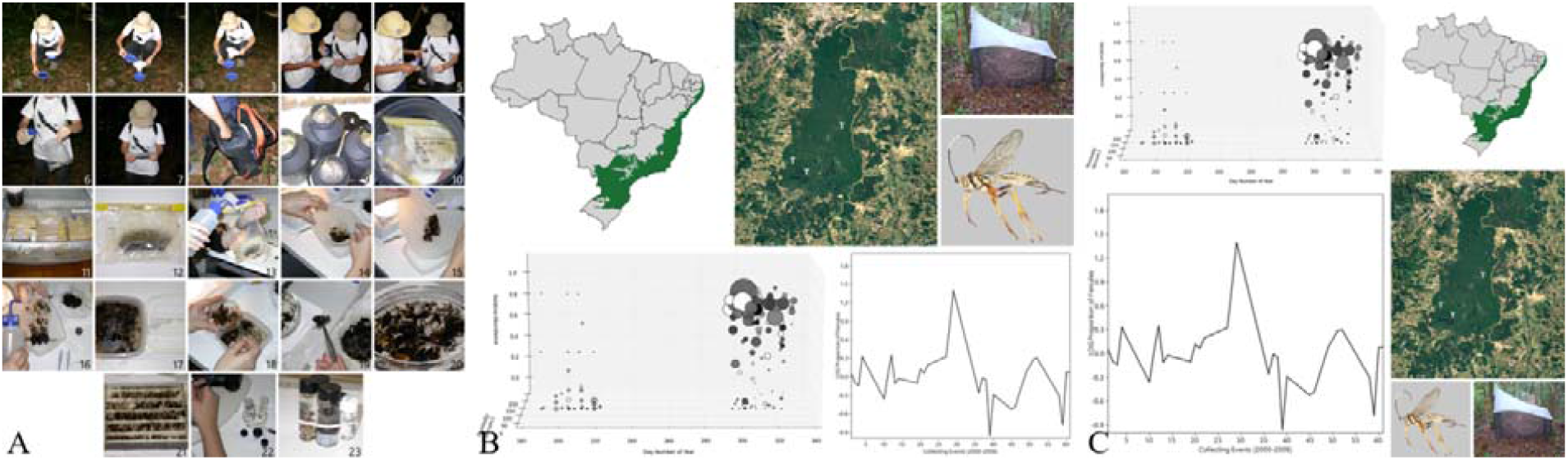
Moericke trap processing, storage, and sorting, and abundance of Cryptini (Ichneumonidae) in a 10-year study (original). **A**, Instant multi panel composition with numbers and automatic font color selection, obtained simply by loading the images (default). **B–C**, Structural clockwise rotation of the layout in 90° increments using button 13.

First, plate construction can be further streamlined by allowing basic operations to be applied simultaneously to multiple selected images, including movement, rotation, flipping, and deletion. Cropping, however, is more appropriately restricted to individual panels to preserve precise control over the resulting image.

User-defined *spacers* (Figure 4) provide convenient, controlled blank areas between images, facilitating balanced compositions with an odd number of panels and avoiding disproportionate panel sizes in the final figure. In Griphus, spacers can be generated by reproducing the proportions of an existing image with a right mouse click or by specifying a width-to-height ratio in the command field.

**FIGURE 4.**
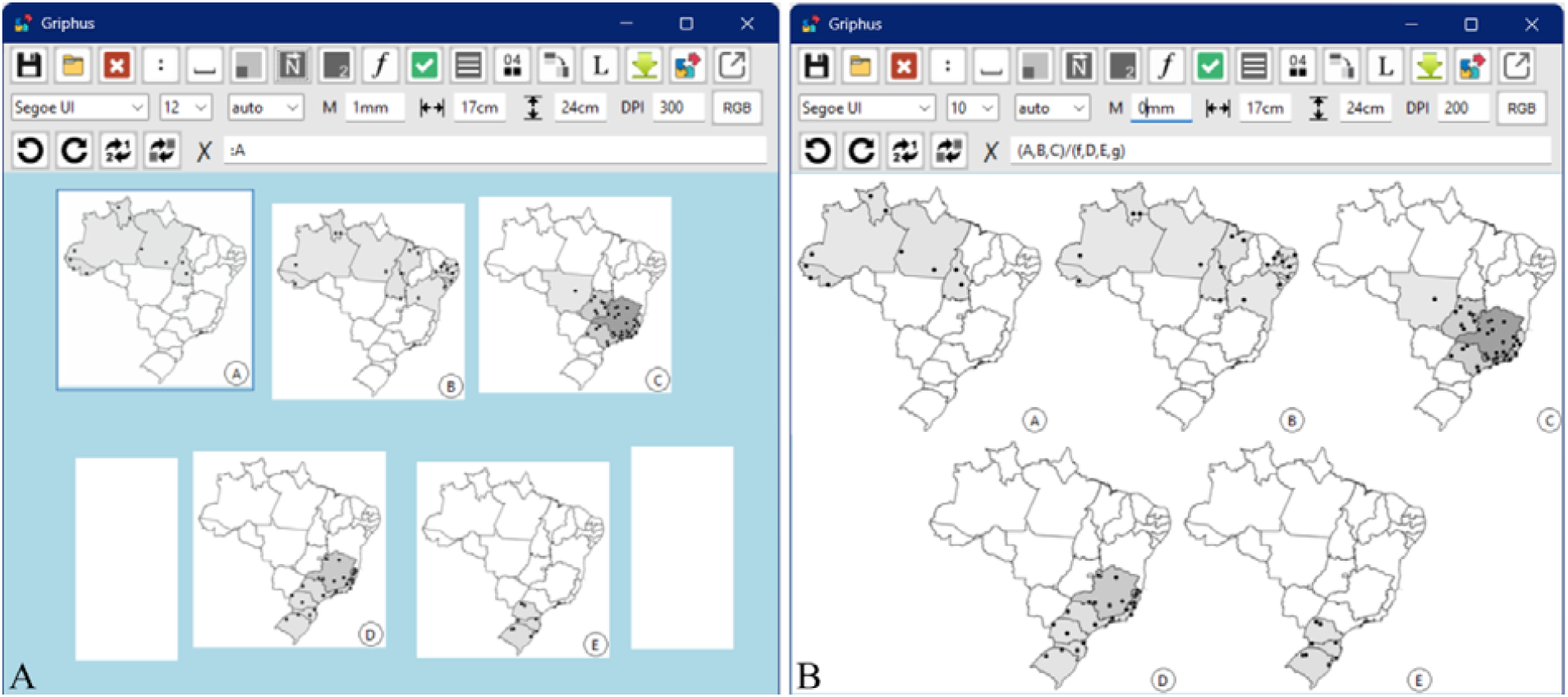
Use of *spacers* (white panels) to align an odd number of species distribution maps (Brazil). **A**, Five square images and two spacers. The first spacer was generated by entering 1:2 in the command line and pressing Enter. The second was replicated with a right mouse click. **B**, Final image generated with button 10.

Aspect-ratio control is also important for producing precisely cropped images, particularly when constructing symmetrical or otherwise consistently organized layouts, and should therefore be readily accessible. In Griphus, a dedicated function calculates and displays the simplest integer ratio for a selected image or symmetrically crops multiple selected images to match the most nearly square aspect ratio among them. A specific ratio can also be supplied through the command field, including ratios expressed as decimal values.

Panel numbering can also be time-consuming and prone to errors, as images must otherwise be labeled individually and in the correct sequence. An evident requirement is therefore to automate panel numbering whenever possible. A simple spatial interpretation is broadly applicable and sufficiently accurate for most layouts: panels can be numbered from the upper left to the lower right of the canvas, proceeding either by rows or by columns. Allowing the resulting sequence to be edited, for example by swapping labels or values, would further extend this approach to accommodate virtually any required numbering scheme.

Griphus implements these approaches and allows numbering to begin from a user-defined numeric or alphanumeric value (e.g., “A1”, “10b”) entered in the command field. Another common requirement, when a circular white background behind labels is undesirable, is to ensure sufficient contrast between each label and the underlying image. Label position at image corners and label color (black or white) should therefore be configurable, ideally with automatic contrast detection. These options are also available in Griphus.

### Use Cases

#### Photographs

Griphus has been extensively used by the author in the preparation of figures for publications in taxonomy, ecology, methodological, and evolutionary research. Early results can be seen in Aguiar et al. (2010), Aguiar and Santos (2012), and Aguiar (2012). More recent uses appear in Supeleto et al. (2020a/b), Aguiar et al. (2024), and Aguiar et al. (2026).

#### Maps and Plots

The software also facilitates the construction of *small multiples*, a visualization approach in which several maps or plots with identical scale and design are arranged in a grid to enable direct comparison across spatial or categorical variables, as proposed by Bertin (1967, 1977) and popularized by Tufte (1990). This strategy is widely used in cartography and data visualization because repeated comparable panels reveal spatial patterns that are difficult to perceive in a single composite map.

In taxonomic studies, especially revisions, distribution data for multiple species are often presented on a single map using different symbols, producing visually crowded figures. A clearer alternative is to use small multiples with one map per species, which can also be ordered from the most northwestern to the most southeastern distribution, as in Figure 3. The program has been used by the author to implement this approach in several publications, including Supeleto and Aguiar (2022) and Supeleto et al. (2022). Griphus includes a preliminary algorithm that arranges loaded maps or plots by visual similarity, which may assist in the automatic organization of small multiples.

## DISCUSSION

The assembly of multi panel figures is currently most commonly performed using general-purpose software or informal techniques. Many online tutorials show how to combine images in presentation software, including widely viewed instructional videos. Published guidelines are also available, such as instructions by PLOS (2019) for combining images in GIMP, and by Nature (2025). However, while informative, these resources provide guidance rather than purpose-built functionality.

General-purpose image editors offer greater flexibility but still require manual coordination of resizing, alignment, spacing, cropping, and labeling. Because these platforms are designed for broad creative work, multi panel construction is possible but requires that users learn and navigate extensive toolsets, which limits iterative testing of multiple, alternative arrangements, discouraging exploratory refinement and optimization.

An early command-line R package by Graumann & Cotton (2018) can generate simple composite panels. Other options are large or highly specialized tools, including FigureJ (Mutter and Zinck 2013) for microscopy data, PyMOL (Chen et al. 2024) for molecular figures, hyve (Ciric et al. 2024) for neuroimaging data, DeTikZify (Belouadi et al. 2024) for TikZ graphics, and EasyFigAssembler (Long and Thu 2026) for omics-related visualizations.

The closest conceptual alternative to the ideas presented herein appears to be PyFigures (Aigouy and Prud’homme 2024). Installation, however, requires familiarity with development environments, and the interface exposes about 140 user-side options distributed across menus, settings, and parameters. In contrast, Griphus works as a standalone application and is intentionally small and focused, with about 40 user-side actions, most accessible within one or two steps, many of which are unavailable in PyFigures. Griphus requires only a few button clicks from image loading to figure export. Despite its simplicity, key features appear to be unique to Griphus, including plate structure inference from image distribution, automatic numbering functions, scale bar functions, spacers, formula-driven layout, insets, and others.

While the proposed ideas and approaches can be replicated and possibly improved using LLMs, there remains a substantial gap between knowing what should be implemented and producing an organized, reliable, and usable software. LLM-assisted programming does not eliminate this difficulty and remains limited when treated merely as a means of generating one-off solutions. As the codebase grows and requires a more robust architecture, it can quickly become complex, inconsistent, or fragile unless its development is actively directed, systematically reviewed, and repeatedly tested.

Artificial intelligence resources can also perform image compositions directly, but access is currently limited for ample scientific experimentation, particularly with large images and more complex layouts. Precise spatial organization may require highly specific prompting, while remote image processing can be computationally demanding and constrained by platform limitations and subscription quotas, reducing flexibility for rapid iterative experimentation.

### Software Limitations

Griphus is designed as an experimental tool for assembling illustration plates in taxonomic work. Annotation beyond image numbering and scale bars remains intentionally limited. Advanced image transformations and detailed editing should be performed using dedicated image editing software such as GIMP, Photopea, Pixlr or equivalent. The current implementation supports only raster image inputs, which may limit use cases requiring vector graphics such as SVG or PDF. Rotations are limited to 90-degree increments. Automatic layout inference may not always capture all intended semantic groupings and may require manual refinement through formula editing.

### Future Work

Future developments are planned in areas such as improved automatic legend and scale-bar generation, layout suggestions, and the grouping of similar images, with Griphus serving as a testbed for new ideas and approaches. Depending on demand, future work could include a fully browser-based local version and a public release of the software on GitHub. The guiding principle for the software itself, however, is to keep it small and focused. Replicating capabilities already available in general-purpose graphics applications is not an objective, as this would add unnecessary complexity and risk obscuring the novel approaches proposed specifically for multi-panel composition.

## CONCLUSION

The proposed concepts and approaches, together with their implementation in Griphus, formalize multi-panel figure composition as a simple but reproducible and parameterized computational process. By combining direct structural manipulation, layout formalization, and session persistence, they accelerate figure assembly while expanding opportunities for layout experimentation and controlled production of publication-quality figures. Several approaches were specifically conceived and integrated for multi-panel composition and taxonomic work, including spatial interpretation of intended layouts, automatic generation of figure legends, automated scale-bar calculation and reporting, and several other procedures tailored for taxonomic work. Together, these approaches provide a substantially new way of designing and producing plates while maintaining strict control over parameters such as dimensions, spacing, and resolution. Griphus provides a practical implementation in which these ideas can be applied, tested, and further developed. As figure assembly remains an important but often time-consuming stage of scientific publication, the ideas presented here offer a means of improving multi-panel figure production while substantially reducing repetitive manual and intellectual work.

## SOFTWARE AVAILABILITY

Distribution details, documentation, and repository information are available at https://www.systaxon.ufes.br/gri.

## ACKNOWLEDGMENTS

OpenAI is acknowledged for providing initial free trial access to *Codex*, which was instrumental in demonstrating that the tool could be effectively used to complete the development of the software and prepare it for release.

## Notes

https://www.systaxon.ufes.br/gri

https://www.systaxon.ufes.br/gri/Manual.html

### Competing Interest Statement

The authors have declared no competing interest.

### Summary of Updates

The manuscript was substantially revised to emphasize the general problems involved in multi-panel figure composition and the new concepts and approaches proposed to address them, shifting the focus from the software to the new ideas and approaches. The presentation was reframed so that these methodological contributions are introduced first and Griphus is presented as their practical software implementation. The Introduction, Methods, Results, Abstract, and Conclusion were revised accordingly. The role of AI-assisted software development was clarified, including the distinction between the original concepts and approaches developed by the author and the use of OpenAI Codex for their implementation, refactoring, testing, and development into distributable software. The description of several approaches and specialized functions for multi-panel composition was expanded and clarified, including layout inference, symbolic layout formulas, automated panel numbering, aspect-ratio control, spacers, scale bars, figure legend generation, physical plate dimensions, resolution control, and direct interface manipulation. Future development and software distribution were also clarified. Figures and figure legends were revised.

